# Characterizing Complex Upper Limb Movements with and without Visual Feedback in Typically Developing Children

**DOI:** 10.1101/2025.03.28.645998

**Authors:** Rachel L. Hawe, Alexandria N. Richardson, T. Lu

## Abstract

The development of upper limb movements has been primarily described through reaching movements, which may not have the complex motor planning and execution demands of many daily tasks. In this study, we introduced a complex task in which individuals had to navigate their hand from a start target through two openings in a simple maze to reach an end target. In half the trials, participants received visual feedback of their hand position, and in half of the trials they did not. Thirty-one participants ages 8 to 17 years completed the study. We found that with visual feedback, reaction time, number of speed peaks, movement time, and hand path length all decreased with age. Number of speed peaks, movement time, and hand path length were all increased without visual feedback, however, the impact of removing visual feedback was greatest on the younger children and decreased with age. Our results demonstrate that complex upper limb movements are refined across childhood and adolescence, with a decreasing reliance on visual feedback likely due to increasing feedforward control and improved ability to use proprioceptive feedback. This task can be applied to clinical populations such as cerebral palsy to assess impairments in motor planning and execution as well as determine how proprioceptive impairments contribute to complex movements.

**NEW AND NOTEWORTHY:** This study used a novel paradigm to examine how children plan and execute complex upper limb movements, both when they have visual feedback of hand position available and when they do not. We show that with age, performance with visual feedback improves reflecting advancing motor planning and execution abilities. Older children were less impacted by removing visual feedback than younger children, likely due to improved feedforward control and use of proprioceptive feedback.

## INTRODUCTION

The development of upper limb movements has primarily been described using reaching tasks in which a child moves their hand from one target location to another, with or without a grasping component.^1–6^ While this has provided information on the developmental trajectory of upper limb reaching movements, such reaching tasks do not fully describe the complexity of upper limb movement routinely encountered in daily life.^7^ In many daily tasks, reaches do not simply have to cover a linear distance, but trajectories are constrained by the need to navigate around obstacles such as when reaching around objects on a crowded desk or dinner table, or the need to pick up or place one toy figurine without disrupting the others in the play scene. These higher complexity movements require more advanced motor planning and impose additional accuracy constraints on the entire trajectory rather than the end location.^8^ However, the development of more complex limb movements throughout childhood and adolescence has not been characterized.

Prior research in adults has demonstrated that motor control processes are different when the trajectory of the movement is essential to successfully complete the task rather than when only end point accuracy is required.^9^ When the trajectory must be planned to navigate around obstacles, the reaction time is slower, representing a stage of motor planning when the movement trajectory is explicitly represented, which is not seen in simple point to point movements.^8^ Whether the motor system acts as a trajectory controller may also depend on the explicit instructions and goals of the task. When the goal is simply to reach an end location, deviations from the trajectory do not need to be corrected as long as the hand is moving toward the final goal. However, when tasks require specific trajectory control, such as to navigate around obstacles, the motor system is able to act as a trajectory controller and will return to the initial trajectory following a perturbation.^9^ Since most work on upper limb development has used reaching tasks, the development of trajectory control is not known.

For both simple and complex movements, visual and proprioceptive feedback contribute to the ability to perform accurate movements. In a point to point reaching movement, central vision is typically fixated on the end target, with peripheral vision monitoring the limb position until the hand comes into central vision as it nears the target.^10–13^ Prior work in which the visual feedback during a goal-directed reaching movement is perturbed using a “cursor jump” paradigm has shown that movements are under constant visual monitoring.^14–16^ In order to navigate around obstacles, visual attention must be directed to both the obstacles as well as monitoring the limb position throughout the trajectory. When visual feedback of the limb position is not available, individuals must rely on proprioceptive feedback as well as increased feedforward control.

The way children integrate and weigh visual versus proprioceptive information, and the extent to which they are reliant on visual feedback to guide limb movements changes throughout development. In studies of school-aged children, children generally become more adept at using proprioceptive feedback and less reliant on visual feedback with increasing age.^17–23^ The increased ability to use proprioceptive feedback corresponds with an increase in proprioceptive acuity that occurs through childhood and adolescence.^23–27^ The reliance on visual feedback may also be tied to whether children use feedforward or feedback control, with children generally shifting from feedback to feedforward control with age.^23,28,29^ A limitation of prior work is the simplicity of the movements studied, mostly point to point reaching movements. When performing more complex movements, the reliance on visual feedback of the limb may be greater to maintain the trajectory control, as well as a different need for feedback versus feedforward control processes.

In this study, we introduce a novel paradigm to examine motor planning and execution during complex upper limb movements. In this paradigm, children must move their hand through a simplistic “maze” to move from a start target to an end target. The nature of the maze requires children to both plan and execute a specific trajectory rather than just a point-to-point reach. Motor planning processes can be examined in both the reaction time, as well as the trajectory and number of speed peaks. We would expect that if children plan the entire trajectory prior to initiating movement, their pathlength would be shorter reflecting that they have found the shortest path through, and they would have fewer corrective movements represented by the number of speed peaks. This paradigm also allows us to examine the role of visual feedback of the hand position. By manipulating whether children have visual feedback of their hand, we can determine their reliance on visual feedback to perform the movement. The objectives of this study were to determine both how performance on a complex task change across development, and to determine the reliance on visual feedback of the hand position. We hypothesized that children would improve with age both when visual feedback was available and when it was removed, demonstrating that older children are less reliant on visual feedback.

## METHODS

### Participants

Thirty-two participants were recruited from the community and participated in this study, with 31 completing the full protocol and included in the analysis (average age 12.5±3.4 years, 13 females/18 males; 26 right-handed/5 left-handed). Inclusion criteria was: ages 8-17 years, with no history of neurologic or neurodevelopmental diagnosis (including autism spectrum disorder, developmental coordination disorder, ADHD), or orthopedic condition impacting the upper limbs. Handedness was based on which hand the child writes with. This study was approved by the University of Minnesota Institutional Review Board and all participants gave written assent along with parental consent.

### Experimental Setup

Experimental procedures were performed using the Kinarm Exoskeleton (Kinarm, Kingston, Ontario, Canada). Participants were seated in a modified wheelchair with booster seats used as needed on smaller children. The bilateral exoskeletons were adjusted to the participant’s upper limbs and supported the limbs on the horizontal plane, as shown in Figure 1. Participants were positioned in front of a horizontal display that projected task-related visuals onto the same plane as their upper limbs. An opaque screen occluded vision of the participants limbs, however, visual feedback of the fingertip positions could be displayed, as detailed below. An apron was also placed over the participants upper arms to occlude vision of their proximal limbs. Kinematics are recorded at 1000 Hz.

**Figure 1:**
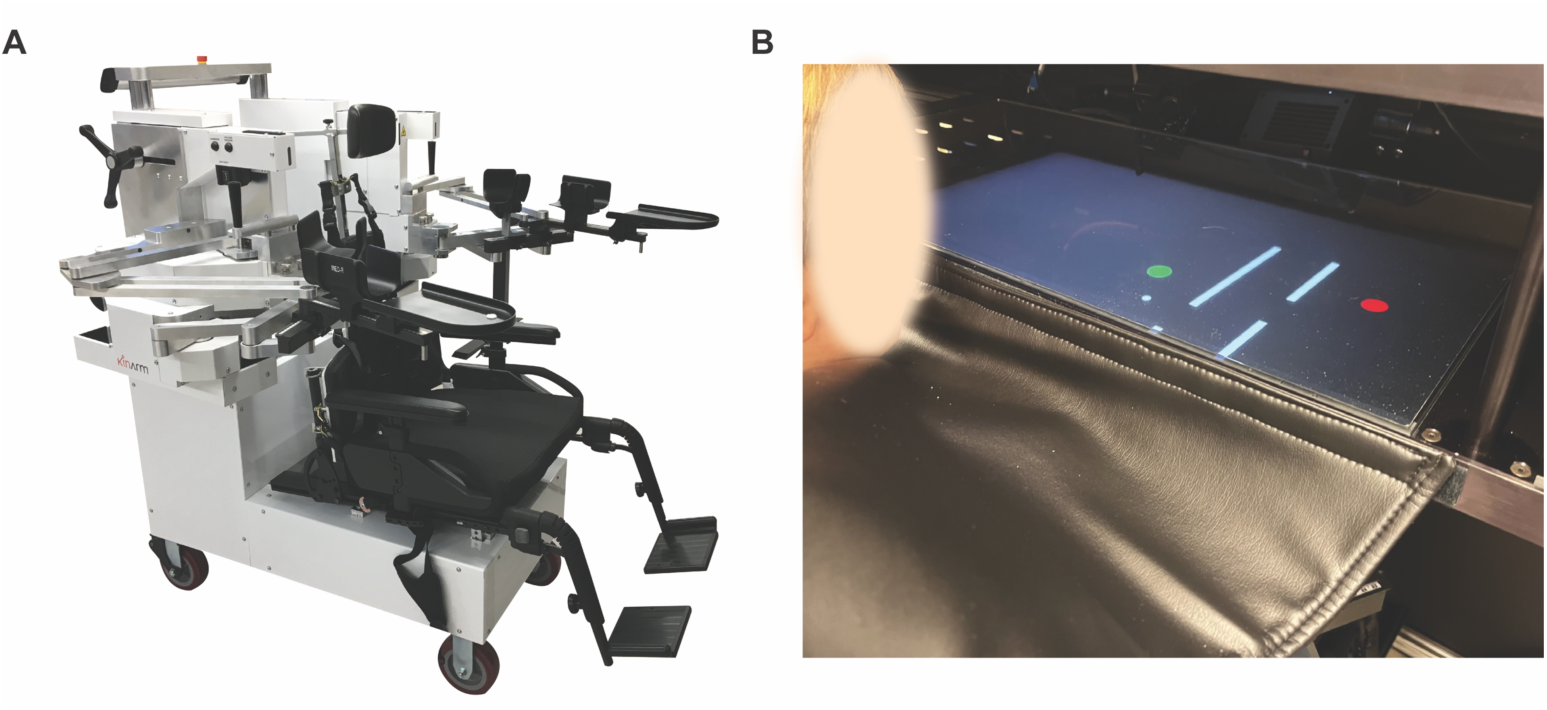
Set-up and task. A) Kinarm exoskeleton is shown with modified wheelchair base and bilateral upper extremity exoskeletons that are adjustable to fit an individual’s upper limbs. B) A 9 yo participant is shown setup in the Kinarm. They are positioned in front of the horizontal screen while their arms are supported by the exoskeleton. The black apron blocks the view of his proximal limbs. The maze is shown on the screen, with the green start target and red final target, and the two bars and passages of the maze. The white dot represents the position of his index fingertip, which is present for trials with visual feedback.

### Experimental Protocol

The task began with participants reaching their hand to a red circular “start target” with a radius of 1.5 cm. In all trials, participants initially received visual feedback of their hand position in the form of a white circular cursor located at the position of the tip of their index finger. Once participants reached the start target, the target changed from red to green. After a random delay interval between 1500 and 3000ms, a maze-like design with two openings and a circular end target (1.5 cm radius) appeared, as shown in Figure 1. In the trials without visual feedback, the feedback of the hand position would also turn off at this point. In the trials with visual feedback, the feedback remained on in the form of the white cursor. Participants were instructed to move their hand through the maze to reach the final target as quickly as possible, without hitting the walls of the maze. Four different configurations of the maze were used, as shown in Figure 2. The horizontal spacing between the walls was 7.4 cm and the vertical openings that the hands passed through were 6 cm. Prior to beginning data collection, the maze was shifted in space to ensure that participants would be able to reach regardless of limb length. If participants contacted the walls of the maze, which was considered an error, the walls would turn red for the duration that the hand was in contact with it. When participants reached the end target, the target would turn from green to red. Participants completed each of the four maze variation 6 times, 3 times with visual feedback and 3 times without. The maze variations and visual feedback conditions were randomized for 24 total trials for each hand. Participants completed the task for both their dominant and non-dominant hand. Mazes were flipped for the right and left hand (right hand version shown in Figure 2). The right hand progressed from the left side to the right side, while the left hand progressed from the right side to the left side.

**Figure 2:**
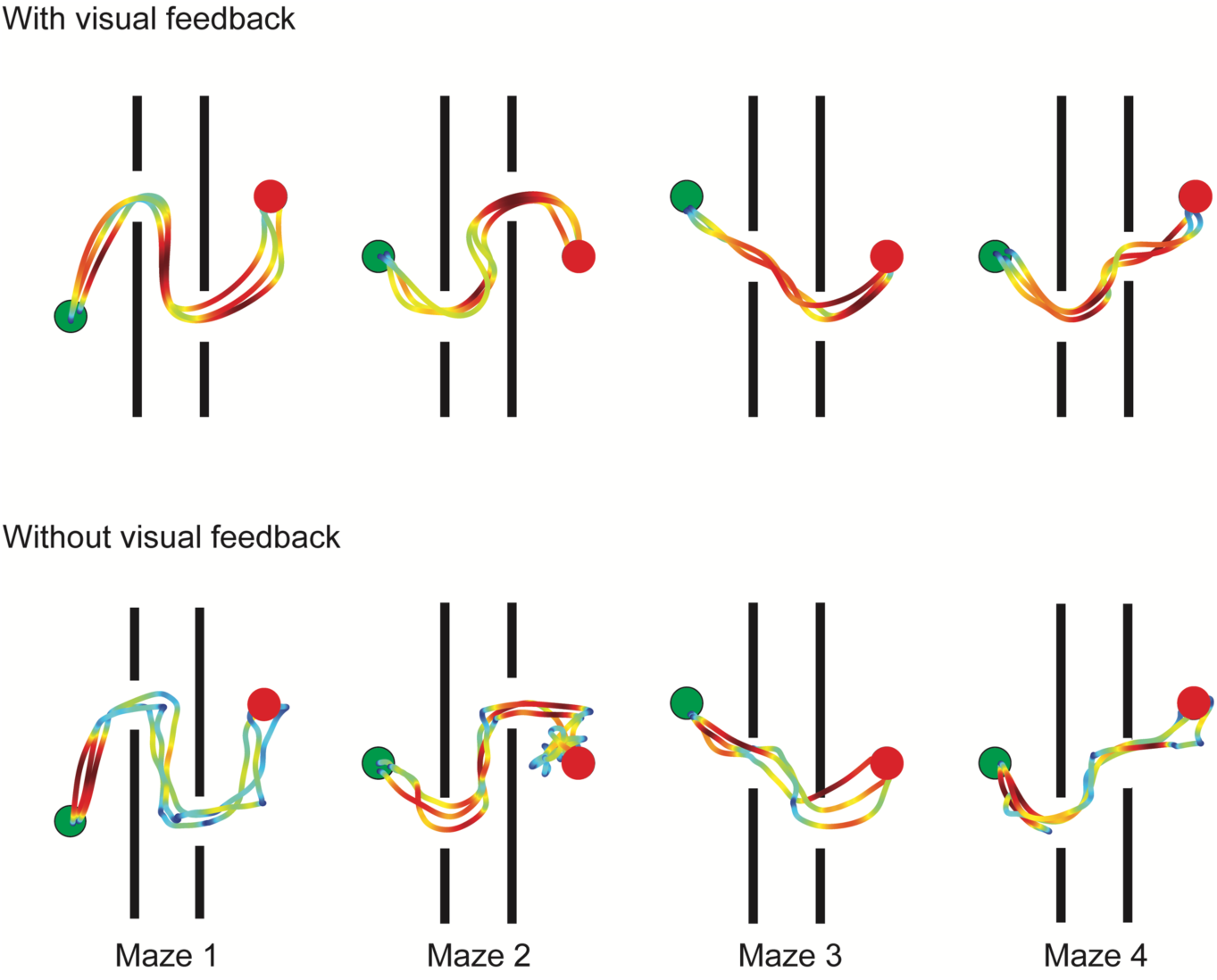
Maze designs and example trajectory data. The four different maze designs can be seen (same designs for with and without visual feedback conditions), including start (green circles) and end target locations (red circles). These mazes would be for the right hand, and the mazes would be flipped for the left hand. The trajectories of the hand are shown for each trial for a 10 year old participant. The trajectories show the hand position from when the maze turned on to when they reached the end target. The color of the trajectory represents the speed of the hand at that position, with warmer colors (red) indicating faster speeds and colder colors (blue) indicating slower speeds, as normalized for each individual trial. Each maze shows the three different trials.

### Analysis

All analysis was done in Matlab 2022a. Kinematic data collected from the Kinarm was filtered using a sixth order double-pass filter with a 10 Hz cutoff frequency.

Each trial was analyzed from the time period between when the maze and end target turned on until the hand reached the end target. In the case where the participant reached the end target and then left it before returning to it, the last occurrence of reaching the end target was used. For each trial, we derived several measures of spatial and temporal performance.

#### Number of Errors

An error indicates any time the participant hit the walls of the maze. Errors were also further subdivided into number of times they hit the first wall and number of times they hit the second wall to determine how errors may increase across the movement. Participants may have hit the same wall multiple times in the same trial. This was counted as only one error as additional errors on the same wall were typically the child attempting to correct based on the feedback of the wall changing color. Therefore the maximum number of errors in a trial was 2 if they hit both the first and second wall one or more times each.

#### Reaction Time (ms)

To calculate reaction time, we first determined *movement onset* as the last time in which the velocity of the hand exceeded 8% of the maximum velocity, prior to the hand exiting the start target. If, on rare occasions, the velocity did not exceed 8% of the maximum velocity prior to leaving the start target, movement onset was considered to have occurred when the hand left the target. Reaction time was calculated as the time between when the maze and end targets turned on and movement onset. Reaction time is a measure of motor planning.

#### Number of Speed Peaks

The number of hand speed local maxima, as a measure of feedback control, as greater number of speed peaks indicate more corrective movements.

#### Movement Time (ms)

The movement time is the time from movement onset to when the hand reaches the end target. If participants reached the end target and then left it before returning to it, the last occurrence of reaching the end target was counted.

#### Hand Path (m)

Hand path length was calculated as the distance the hand traveled from when the maze and end target turned on to when the hand reached the end target for the last time. Hand path can demonstrate the efficiency of the movement and the amount of advanced planning, as shorter hand paths can be executed if planning for the location of the next passage or end target. Additionally, hand path will be longer with more corrective movements.

### Statistical Analysis

We used linear mixed effects models with random intercepts to examine the effects of age, hand (dominant vs. non-dominant), and visual feedback condition, while controlling for the four different maze configurations. Data from all 48 trials from each participant, rather than averages, were used for this approach. We examined the fixed effects of age (in years), hand (dominant coded as 1, non-dominant as 0), and condition (with visual feedback coded as 1, without visual feedback coded as 0). The maze design (four variations) was included as a random effect. Models initially included interaction terms for age x visual feedback condition and hand x visual feedback condition. If the interaction term was not significant (p>0.05), it was removed from the final model. Separate models were fit for each of the measures (number of errors, reaction time, number of speed peaks, movement time, and hand path. Based on observations that more errors occurred in the second bar than the first, we performed additional linear mixed effects models on the trials without visual feedback only, with fixed effects of bar (bar 1 coded as 1, bar 2 coded as 2), age, hand (dominant coded as 1, non-dominant as 0), and age x bar interaction, with random intercepts for participant and maze. Lasty, we examined correlations with age separately for each hand and visual feedback condition using Pearson correlations. For each variable, the average value across the 12 trials was used in the correlation. For each feedback condition, family-wise error rates were controlled for using the Holm-Bonferroni method (8 correlations for each feedback condition (reaction time, number of speed peaks, movement time, hand path length for dominant and non-dominant arms)).

## RESULTS

Example trajectories color coded for speed are shown in Figure 2. The effects of age, visual feedback condition, and hand (dominant or non-dominant) are described for each measure below, with results of the linear mixed effects models examining the effects of age, visual feedback condition, and hand are shown in Table 1.

**Table 1:** Results of Linear Mixed Effects Models. Bolded values indicate significant term.

|  | **Age (years)**<br>Estimate (SE),<br>p-value | **Hand (1=dom; 0**<br>**= non-dom)**<br>Estimate (SE), p-value | **Visual**<br>**Feedback**<br>**Condition (0 =**<br>**no vis fdbk; 1 =**<br>**vis fdbk)**<br>Estimate (SE),<br>pvalue | **Age × Visual**<br>**Feedback**<br>**Condition**<br>Estimate (SE),<br>p-value | **Hand ×**<br>**Visual**<br>**Feedback**<br>**Condition**<br>Estimate<br>(SE), p-value |
| --- | --- | --- | --- | --- | --- |
| Number<br>Errors | 0.016 (0.009)<br>*p*=0.080 | **-0.047 (0.021)**<br>*p*=0.027 | -0.158 (0.082)<br>*p*=0.054 | **-0.019 (0.006)**<br>*p*=0.002 | -- |
| Reaction<br>Time | **-22.518 (3.356)**<br>*p*=2.78x10<sup>-11</sup> | **-14.301 (6.43)**<br>*p*=0.026 | 11.592 (6.43)<br>*p*=0.072 | -- | -- |
| Number of<br>Speed Peaks | **-0.399 (0.120)**<br>*p*=0.001 | -0.278 (0.200)<br>*p*=0.164 | **-4.95 (0.770)**<br>*p*=1.76x10<sup>-10</sup> | **0.117 (0.056)**<br>*p*=0.049 | -- |
| Movement<br>Time (ms) | -99.3 (56.2)<br>*p*=0.078 | **-226.2 (72.6)**<br>*p*=0.002 | **-1771.5 (72.6)**<br>*p*=1.05x10<sup>-110</sup> | -- | -- |
| Hand path<br>Length (m) | **-0.009 (0.002)**<br>*p* = 5.63x10<sup>-6</sup> | **-0.039 (0.007)**<br>*p*=2.16x10<sup>-7</sup> | **-0.143 (0.021)**<br>*p*=1.47x10<sup>-11</sup> | **0.004 (0.002)**<br>*p*=0.007 | **0.03 (0.01)**<br>*p*=0.002 |

### Number of errors

Overall, we found errors to be exceedingly rare with visual feedback of the hand position, with only 15 errors occurring across all trials and all participants. Without visual feedback, there was a total of 312 errors across all participants’ trials. The results of the linear mixed effects model (Table 1) show that the fixed effects of age and visual feedback condition were not significant but there was an age × visual feedback interaction. Interpreting this interaction term shows that with visual feedback, the number of errors decreased with age, however, no age-related changes were seen without visual feedback. The number of errors was also less with the dominant hand. Note that due to the large number of overlapping points at 0 or 1 error, we have not shown error as we have with the other measures in Figures 4-7.

We also found that without visual feedback, more errors were made later in the movement trajectory, indicated by more frequently contacting the second bar than the first, as can be seen in Figure 3. Without visual feedback, 62.7% of errors with the dominant arm and 64.7% of errors with the non-dominant arm involved the second wall Linear mixed effects models showed a significant effect of bar (estimate: 0.291, standard error 0.074, p=9.36×10^−5^) and an age × bar interaction (estimate: −0.015, standard error 0.006, p=0.008), indicating that more errors were made with the second bar, but this effect decreased with older ages.

**Figure 3:**
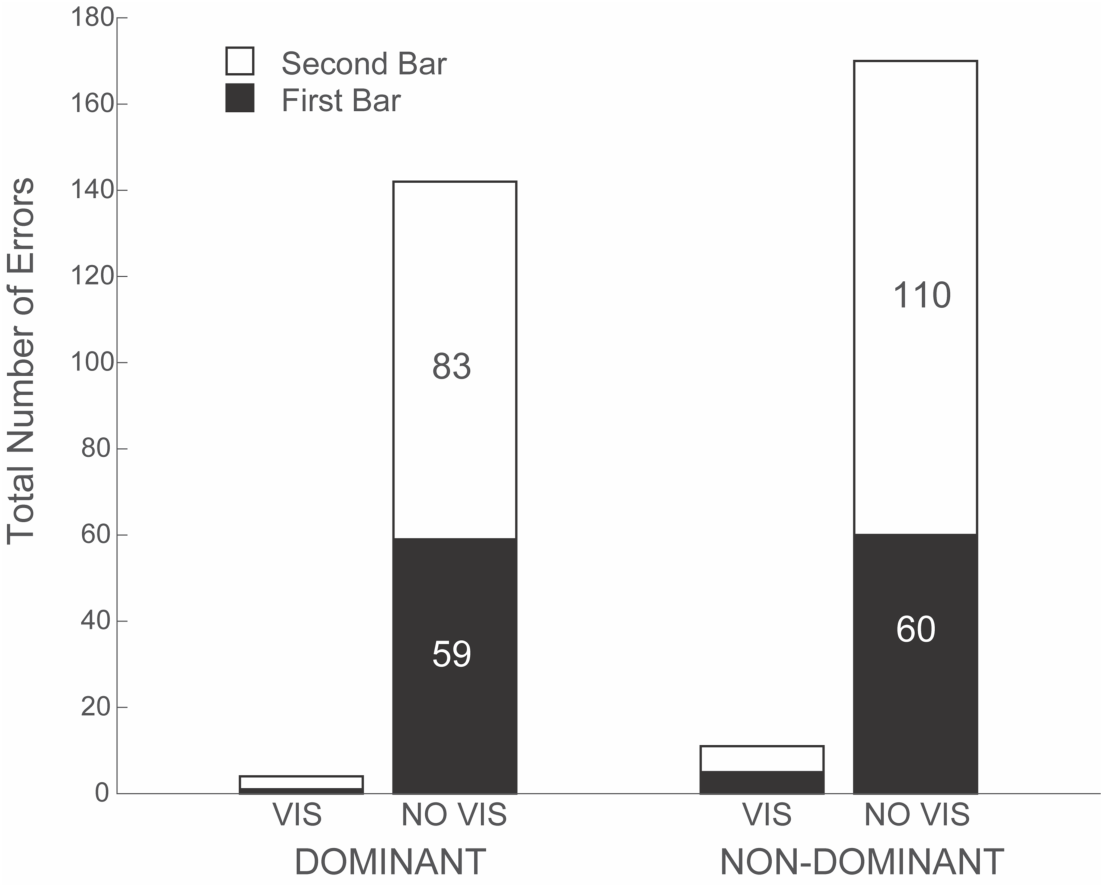
Total errors. The bar graphs display the total number of errors across all participants. The black regions indicate the error was with the first bar, and the white indicates the second bar. An error is considered to have occurred if a participant contacts the “walls” of the maze instead of passing through the passage. There is a maximum of two errors per trial-one with each bar. “VIS” indicates trials with visual feedback of the hand and “NO VIS” indicates trials without visual feedback of the hand.

### Reaction Time

Reaction time results are shown in Figure 4. The linear mixed effects model (Table 1) showed significant fixed effects of age and hand, with reaction time decreasing with age and being shorter with the dominant hand. The lack of change in reaction time between the visual feedback conditions can be appreciated in Figure 4 A and B. Figure 4 C-F also shows the strong correlations between age and reaction time for both hands and both visual feedback conditions.

**Figure 4:**
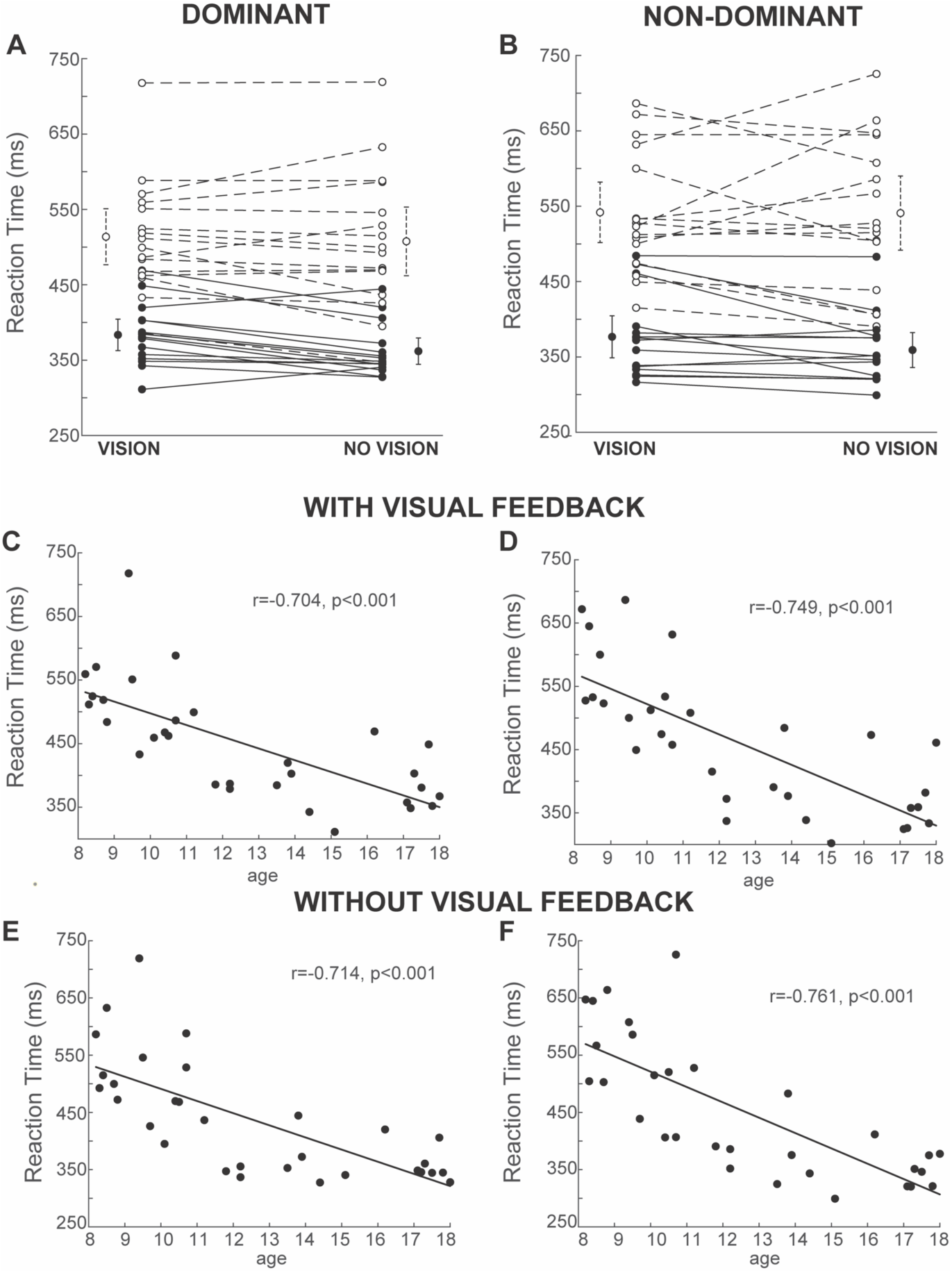
Reaction time. A and B depict the reaction times for each participant for the conditions with visual feedback (“VISION”) and without visual feedback (“NO VISION”) for the dominant (A) and non-dominant (B) hands. The circles indicate the reaction times for each condition, averaged across the three trials, with the line connecting the two conditions. For individuals less than 12 years of age (n=16), data is shown with open circles and dashed lines. For individuals 12 years and older (n=15), data is shown with closed circles and solid lines. Adjacent to the individual data, the group means (<12 years and ≥12 years) are shown with 95% confidence intervals. C-F show the correlation between reaction time and age for the dominant hand (C and E) and non-dominant hand (D and F), and the conditions with visual feedback (C and D) and without visual feedback (E and F).

### Number of Speed Peaks

Fluctuations in speed can first be appreciated in Figure 2, where the hand paths are color coded for relative speed of that trial. With visual feedback, it can be observed that speed is generally modulated (slowed) as the hand moves toward and through the passages and as it reaches the end target. Without visual feedback, there are more non-systematic fluctuations in hand speed showing more corrective and segmented movements. The results of the linear mixed effects model (Table 1) demonstrate the number of speed peaks had significant effects of age, visual feedback condition and age × visual feedback condition interaction. Interpreting the linear mixed effects coefficients, we find that there are approximately 5 more speed peaks without visual feedback of the hand compared to with visual feedback, which can be appreciated in Figure 5 A and B. The age x visual feedback condition interaction demonstrates that the impact of visual feedback was greater on younger children. Correlations with age (Figure 5 C-F) show significant correlations for the with visual feedback condition, but with Holm-Bonferroni corrections, the correlations were not significant without visual feedback.

**Figure 5:**
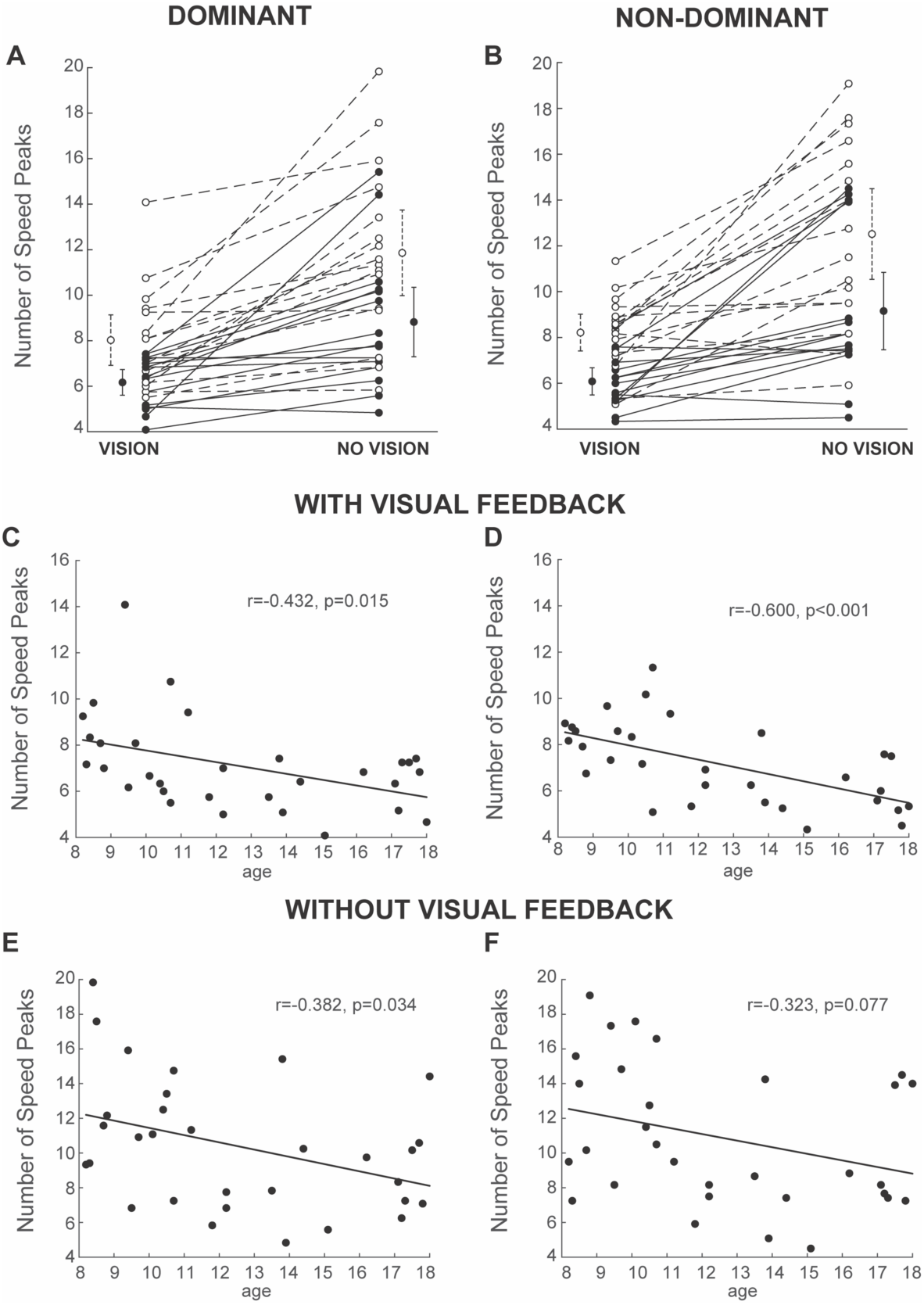
Number of speed peaks. A and B depict the number of speed peaks for each participant for the conditions with visual feedback (“VISION”) and without visual feedback (“NO VISION”) for the dominant (A) and non-dominant (B) hands. For individuals less than 12 years of age (n=16), data is shown with open circles and dashed lines. For individuals 12 years and older (n=15), data is shown with closed circles and solid lines. Adjacent to the individual data, the group means (<12 years and ³12 years) are shown with 95% confidence intervals. C-F show the correlation between number of speed peaks and age for the dominant hand (C and E) and non-dominant hand (D and F), and the conditions with visual feedback (C and D) and without visual feedback (E and F).

### Movement Time

The linear mixed effects model (Table 1) found significant main effects of hand and visual feedback condition. Movement time was 226 ms faster with the dominant hand, and 1771 ms faster with visual feedback. The difference between with and without visual feedback can be appreciated in Figure 6 (A and B). Correlations with age for each hand and each visual feedback condition using Bonferroni-Holm corrections showed significant correlations for each hand with visual feedback but not without visual feedback (Figure 6 C-F).

**Figure 6:**
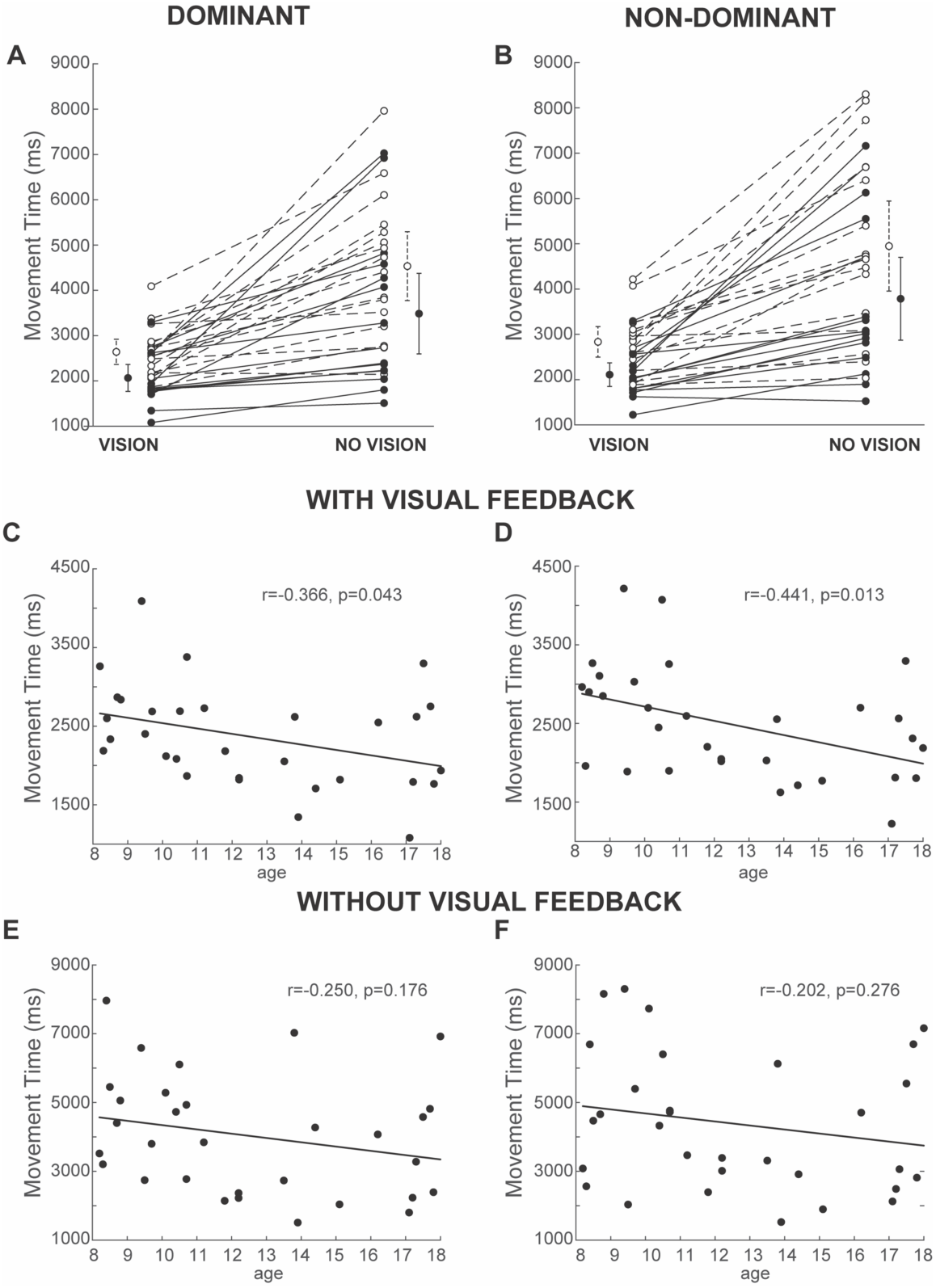
Movement time. A and B depict the movement times for each participant for the conditions with visual feedback (“VISION”) and without visual feedback (“NO VISION”) for the dominant (A) and non-dominant (B) hands. The circles indicate the movement times for each condition, averaged across the three trials, with the line connecting the two conditions. For individuals less than 12 years of age (n=16), data is shown with open circles and dashed lines. For individuals 12 years and older (n=15), data is shown with closed circles and solid lines. Adjacent to the individual data, the group means (<12 years and ³12 years) are shown with 95% confidence intervals. C-F show the correlation between movement time and age for the dominant hand (C and E) and non-dominant hand (D and F), and the conditions with visual feedback (C and D) and without visual feedback (E and F).

### Hand path length

The hand paths with visual feedback and without for a participant can be seen in Figure 2. As shown in Table 1, hand path length had significant fixed effects of age, hand, and visual feedback condition, as well as age x visual feedback condition and hand x visual feedback condition interactions. Hand path was shorter with visual feedback, with the presence of visual feedback having a greater impact on the non-dominant hand and for younger children. The differences between the visual feedback conditions can be seen in Figure 7 A and B. Hand path was also shorter with the dominant hand. The decrease in hand path with age with visual feedback can be seen in Figure 7 C and D. With Holm-Bonferroni corrections, the correlations between age and hand path were not significant without visual feedback (Figure 7 E and F).

**Figure 7:**
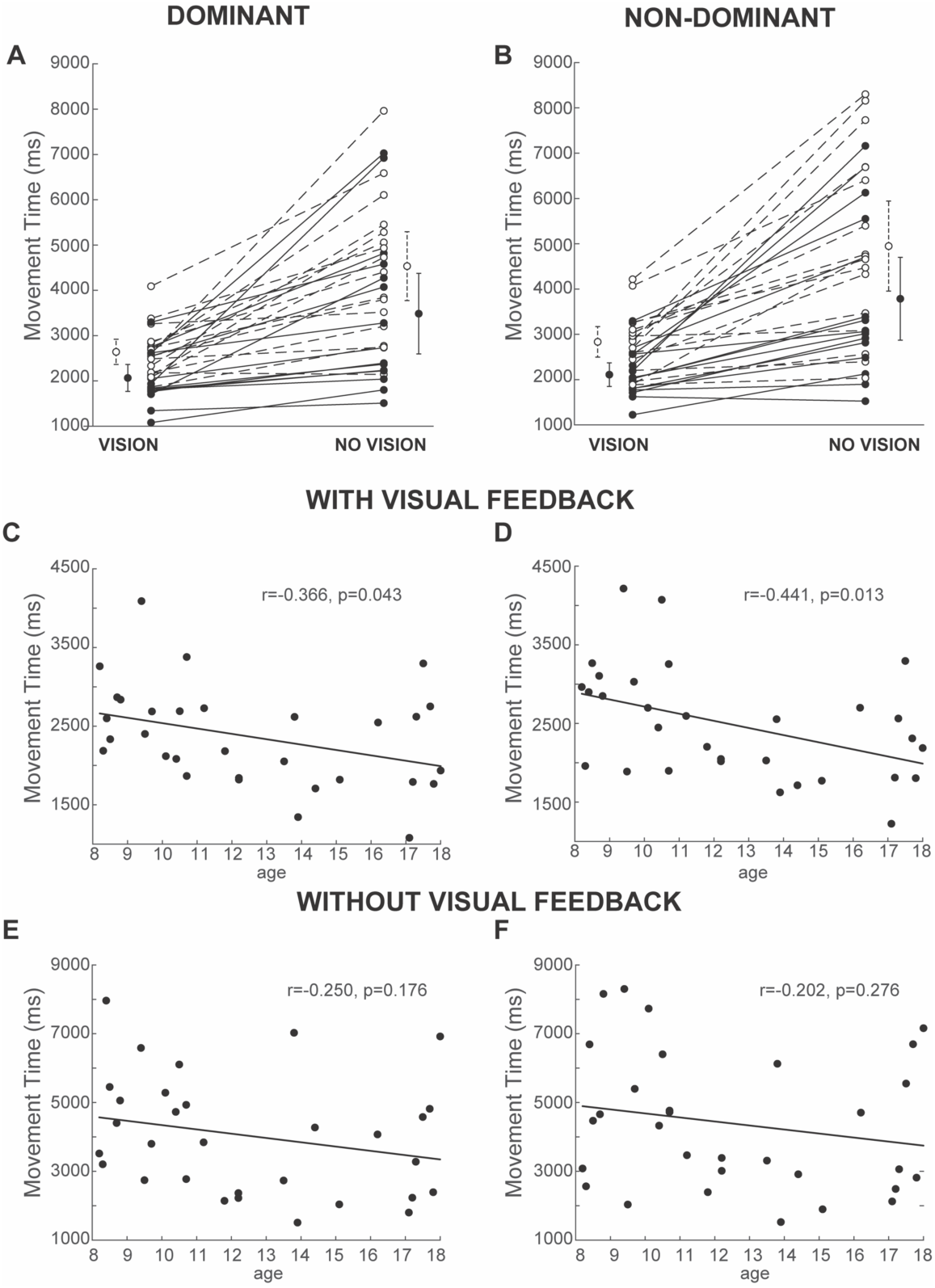
Hand path length. A and B depict the hand path lengths for each participant for the conditions with visual feedback (“VISION”) and without visual feedback (“NO VISION”) for the dominant (A) and non-dominant (B) hands. The circles indicate the path lengths for each condition, averaged across the three trials, with the line connecting the two conditions. For individuals less than 12 years of age (n=16), data is shown with open circles and dashed lines. For individuals 12 years and older (n=15), data is shown with closed circles and solid lines. Adjacent to the individual data, the group means (<12 years and ≥12 years) are shown with 95% confidence intervals. C-F show the correlation between path length and age for the dominant hand (C and E) and nondominant hand (D and F), and the conditions with visual feedback (C and D) and without visual feedback (E and F).

## DISCUSSION

Our results characterize the development of motor planning and execution for a complex upper limb task both with visual feedback of the hand position and without visual feedback. With visual feedback, we found improvements with age in decreasing reaction time, fewer speed peaks, shorter movement times, and shorter hand path lengths. Removing visual feedback impacted all parameters except for reaction time. We found that the effect of removing vision on the number of errors, number of speed peaks, and hand path length was greater for younger children than older children. We found that without visual feedback, errors were more likely to occur later in the trajectory.

The motor planning required for this task is more complex than for a simple reaching task. As there is not just a single target to reach to, individuals must instead interpret the visual scene of the maze and then plan their trajectory to navigate the maze and reach the end target. Prior work has demonstrated that in adults, when a specific trajectory is required to successfully complete the task, there is a distinct planning phase where the trajectory is mapped out.^8^ To our knowledge, this is the first time examining the planning of complex trajectories in children. Several results relating to performance with visual feedback point to advancing motor planning abilities with increasing age. Reaction times significantly decreased with age, demonstrating more efficient motor planning processes in older children and adolescence. We found the hand path length also decreased with age. This likely indicates that older children had more anticipatory planning of the trajectory, allowing them to take the optimal path based on where their hand needed to go next. For instance, in passing through the first opening, if planning ahead and knowing the next opening was more distal, they could navigate the first opening at the distal end, contributing to a shorter path. Anticipatory planning has commonly been examined with end-state comfort paradigms, which examine whether individuals may initially grasp an object in an uncomfortable or awkward posture in anticipation of a comfortable final posture. An example is grasping an overturned glass with the thumb positioned down (pronated) to then have a comfortable position when the glass it turned upright. While results on the end-state comfort effect vary in children, overall they suggest that anticipatory planning emerges across childhood up to age 12, though the exact age may vary by paradigm and factors such as precision requirements and number of steps.^30^ Another indicator of improved planning was that found that the number of speed peaks decreased with age. The number of speed peaks are representative of corrective movements. With more advanced planning and feedforward control, less corrective movements would be expected.

While reaction time was strongly correlated with age for both visual feedback conditions, there was no impact on visual feedback condition on reaction time. We had hypothesized that reaction times would be longer to represent a shift toward more feedforward control in the absence of visual feedback. Our results instead show that the planning process was the same whether children had visual feedback available or not. Prior work in adults with visual feedback has shown that when the trajectory rather than just an end goal is important for a task, the trajectory is explicitly planned prior to movement onset.^8^ Our results would support this and indicate that the entire trajectory is pre-planned whether or not visual feedback will be available. Specific to children, Bard et al. found that when a reaching task required directional control, reaction time also did not differ between having visual feedback and not, as that direction was programmed regardless.^18^ Our results show that with a more complex task than Bard, the same holds true. It is possible that if our task required greater accuracy constraints throughout the trajectory, such as by having more narrow openings in the maze, we may have seen additional time to plan without visual feedback to account for the accuracy demands. An alternative explanation of the same reaction times is that since the conditions were intermixed in a random order, habit may influence reaction times. Prior work has shown that if the condition where reaction time is prolonged is intermixed with trials where a shorter reaction time would be expected, the shorter reaction times are lengthened as a force of habit.^8,31,32^

The difference between with and without visual feedback decreased with age for number of speed peaks, movement time, and hand path length. This is likely due to two processes. First, as children mature, they develop more feedforward control. Therefore, they are less reliant on vision to make corrections as the movement has been “pre-programmed.” In studies of aiming movements, it has been shown that younger children plan only the initial movement and rely on online feedback during the movement. On the contrary, older children have more feedforward control and rely less on online feedback.^33–35^ The age at which this transitions has varied by study but typically between 9 and 11 years old. However, these studies have used only a simple reaching paradigm. An additional explanation is that proprioception has improved as children develop, and that would contribute to less of an impact on having visual feedback removed. In the future, to distinguish between proprioceptive feedback and improved feedforward control, we could introduce a version of the maze in which the placement of the openings is not known until children begin moving, removing the ability to plan the movement in advance and instead needing to rely on proprioception.

We found that without visual feedback, more errors occurred later in the movement trajectory, as shown by participants more commonly contacting the second bar of the maze rather than the first, though this effect decreased with age. This can give insight into how effectively proprioception can be used to control the trajectory. Limb position drift has been well documented when visual feedback is removed, meaning the location of the limb drifts away from the target locations, with the amount of drift accumulating over repeated trials.^36–38^ Results from deafferented patients has shown that in the absence of both visual and proprioceptive feedback, the amplitude or movement distance is preserved but spatial accuracy decreases across the movement sequence.^39^ Specific to children, Hay et al. has found that without visual feedback, children can be accurate at the beginning of a series of movements, however, error increases over the series of movements.^23^ As younger children have less refined proprioception,^24–27^ it is therefore not surprising that their errors increased as the movement progressed.

There were several limitations of this study. First, children were not specifically instructed on what to do if they made an error. Therefore, some children continued onwards with the maze never passing through the opening, while other attempted to first proceed through the opening correctly, potentially repeatedly contacting the wall. To account for this, we only counted a single error on each wall. Additionally, our participants were not evenly distributed across ages, with less participants in the early teenage years. While this is likely a period of slower development, this may have impacted our results. The fact that this paradigm was done in an augmented reality environment without real consequences of hitting the obstacle may influence performance, as individuals may have prioritized not making an error more if it could result in breaking an item or spilling a glass, for instance, and may have had different movement trajectories.^40,41^

While the present work was in typically developing children, future work will examine performance on this task in clinical populations, namely unilateral cerebral palsy. There is a critical need for assessments that involve more complex movements, as typical reaching paradigms may not be sensitive to individuals with more mild impairments that can still significantly impact performance in daily life. For instance, in a robotic visually guided reaching task, Kuczynski et al. found that even with the sensitivity of robotics, many measures found only a minority of children with unilateral cerebral palsy were impaired compared to a normative population.^42^ This likely does not suggest that the remaining children do not have impairments, but rather the movement required was not complex enough to detect them. Examining the role of visual feedback will also allow for the assessment of proprioception as well as visuomotor integration, both of which are commonly impaired and impact movement.^43,44^ The present work provides the first normative data in typically developing children which will inform our future detection of impairments in clinical populations.

## CONCLUSIONS

In this study, we introduced a novel paradigm to exam motor planning and execution during a complex upper limb task with and without visual feedback. Overall, we found motor planning and execution abilities improved with age with visual feedback. Older children were less impacted by the removal of visual feedback, likely due to improved feedforward control and ability to use proprioceptive feedback. Future work will use this paradigm in clinical populations to assess motor planning and execution in a complex upper limb task.

## DATA AVAILABILITY

Data is available by reasonable request to the corresponding author.

## ACKNOWLEDGEMENTS

This work was funded by an Academy of Pediatric Physical Therapy Research Grant. We would like to thank the children and families who participated.

## AUTHOR CONTRIBUTIONS

RLH conceived and designed research, analyzed data, interpreted results of experiments, prepared figures, drafted manuscript; ANR performed experiments, edited and revised manuscript, approved final version of manuscript; TL performed experiments, approved final version of manuscript.

